# The Personalized Proteome: Comparing Proteogenomics and Open Variant Search Approaches for Single Amino Acid Variant Detection

**DOI:** 10.1101/2020.12.11.419523

**Authors:** Renee Salz, Robbin Bouwmeester, Ralf Gabriels, Sven Degroeve, Lennart Martens, Pieter-Jan Volders, Peter A.C. ’t Hoen

## Abstract

Discovery of variant peptides such as single amino acid variant (SAAV) in shotgun proteomics data is essential for personalized proteomics. Both the resolution of shotgun proteomics methods and the search engines have improved dramatically, allowing for confident identification of SAAV peptides. However, it is not yet known if these methods are truly successful in accurately identifying SAAV peptides without prior genomic information in the search database. We studied this in unprecedented detail by exploiting publicly available long-read RNA seq and shotgun proteomics data from the gold standard reference cell line NA12878. Searching spectra from this cell line with the state-of-the-art open modification search engine *ionbot* against carefully curated search databases resulted in 96.7% false positive SAAVs and an 85% lower true positive rate than searching with peptide search databases that incorporate prior genetic information. While adding genetic variants to the search database remains indispensable for correct peptide identification, inclusion of long-read RNA sequences in the search database contributes only 0.3% new peptide identifications. These findings reveal the differences in SAAV detection that result from various approaches, providing guidance to researchers studying SAAV peptides and developers of peptide spectrum identification tools.

## Introduction

Proteomes display significant inter-individual variability ^1,2^ and personal proteomes may delineate disease risk and pave the way for personalized disease prevention and treatment. Personalized cancer treatment, for instance, is already instigated based on the detection of peptides containing single amino acid variants (SAAVs) that often serve as excellent biomarkers ^3–8^. Detecting these SAAV peptides reliably, however, is a formidable challenge. Previously, scientists looked for protein evidence of a small number of variants in particular and resorted to targeted proteomics approaches such as selected reaction monitoring (SRM) ^9–12^. Alternatively, BLAST-like query tools such as peptimapper and PepQuery ^13,14^ or database tools like XMAn v2 ^15^ and dbSAP ^16^ can be used to investigate single events ^17,18^. Proteogenomics, the integration of genome and transcriptome information, is a more holistic and higher-throughput form of mass spectrometry- (MS-) based detection of variant peptides.

A main limiting factor of SAAV peptide (called ‘variant peptide’ for the remainder of the manuscript) detection with shotgun proteomics is the tandem mass spectrometry (MS/MS) technology itself. Since MS/MS spectra are generally too noisy to call a peptide sequence de novo, current MS/MS analysis methods rely on a database of known peptides. This limits the ability to detect unknown peptides such as variant peptides. The most flexible way to detect variant peptides is an exhaustive search; allowing any possible amino acid substitution at any position in the peptide sequence ^19,20^. However, this strategy increases the search space immensely to a point where it is no longer useful in practice. The larger search space leads to ambiguity in peptide identification and thus a high number of false positive hits ^21,22^. Therefore, more careful curation of sequences in the search database pays off.

Databases of peptides containing variants from dbSNP have been created to facilitate the search for SAAVs ^3,16^, and simply adding these variant peptides to the database showed promise early on ^3,23^. Not all dbSNP variants however, are expected to be found in every sample, and including them all may lead to false identifications ^24^. In addition, rare and unique variants may be overlooked. A proteogenomics approach where only those variant peptides predicted from genome or transcriptome information are added to the peptide search databases, can improve their detection. Proteogenomics pipelines have streamlined this process of incorporating personal genome information into a proteomic search database ^25–29^. In addition, there is evidence that including correct sequence variant information, including often-overlooked sample-specific indels and frameshifts, improves variant peptide identification workflows ^30^. Yet, false discovery rate (FDR) correction is needed to compensate for the increase of database size and complexity ^21,22^. When searching for evidence of specific peptides such as variant peptides, an additional subset specific FDR correction should be made ^31^.

In addition to SAAVs, alternative splicing may also introduce sample specific peptides. Alternative splicing is commonplace as 90% of genes undergo alternative splicing ^32^. Since protein reference databases do not cover all protein isoforms produced by alternative splicing, sample-specific transcriptome information is advantageous. Typically, the information on alternatively spliced sequences comes from RNA sequencing. Short-read RNA seq is however not ideal for properly capturing the complete splicing patterns and the resulting open reading frames (ORFs). Traditionally, this is circumvented by including 3- or 6-frame translations of the sample’s transcriptome. However, this approach was found to expand the database far too much for eukaryotic organisms, leaving few remaining hits after FDR correction ^33^. Studies utilizing long-read RNA seq frequently discover previously unannotated transcript structures. Thus, full-length transcripts may add essential information for correct ORF prediction and peptide identification.

An emerging alternative to proteogenomics methods for the detection of variant peptides is the ‘open search’ method. This allows unexpected post-translational modifications and amino acid substitutions in the peptide spectrum match, while maintaining accurate FDR and a workable computation time. Using sequence tag-based approaches, the search space is narrowed with de novo sequence tags, which makes room for the addition of all possible SAAV peptides in the search space ^34–38^. These methods were historically not as effective as classical proteogenomics searches in finding variant peptides, since there is difficulty in discerning between post-translationally modified and SAAV peptides. However, this situation has recently improved with the inclusion of optimized probabilistic models ^39^. One implementation of the tag-based method improved with such models is *ionbot* (manuscript in preparation; compomics.com/ionbot), which is a machine learning search engine that uses MS2PIP ^40^ and ReSCore ^41^ to significantly improve the accuracy of peptide match scoring.

The main objective of this study is to compare a previously established proteogenomics approach based on long-read sequencing with a recently-developed open search method for the detection of true variant peptides. In simpler terms, we compare a genome-informed search space with typical spectrum identification settings to a genome-uninformed search space with advanced identification settings. We aim to understand the power of, and potential biases associated with, using an open search method without prior information about the genome. For this, we make use of high-confidence nucleotide sequencing and (ultra)-deep proteomics data from a gold standard cell line NA12878. Using correct ORFs from the long-read transcriptome and high-confidence phased variants belonging to this cell line, we gain a unique perspective on exactly what advantages can be gained by each approach.

## Experimental section

### NA12878 Data sources

Variant information was obtained from Illumina platinum genomes (ftp://platgene_ro@ussd-ftp.illumina.com/2017-1.0/hg38/small_variants/NA12878/). The reference genome used was GRCh38, which can be downloaded from the pre-computed 1000 genomes GRCh38 BWA database at ftp://ftp-trace.ncbi.nih.gov/1000genomes/ftp/technical/reference/GRCh38_reference_genome/ (with decoys). Transcript structures for NA12878 were sourced from the ONT consortium ^42^. The reference transcriptome and proteome are from GENCODE v29.

Shotgun proteomics data came from the ^43^ study, downloaded from Peptide Atlas (http://www.peptideatlas.org/PASS/PASS00230). This dataset consists of 417 TMT6plex runs from 54 samples, with the reference tag (126.77) on NA12878 in every case.

### Creation of the search databases

Two sets of two search databases were created from the two sources of sequences: the ONT transcriptome and GENCODE-predicted coding transcriptome sequences. The first set contained sequences of each of the sources individually (referred to as ‘ONT’ or ‘Ref’). The second set consisted of the union of the two sources, one database with NA12878-specific variants included (variant-containing, VC) and one database without (variant-free, VF). A simple depiction can be found in Figure 1A, while the detailed full workflow can be found in Figure S1. Each database had MaxQuant ^44^ contaminant sequences appended before search.

**Figure 1.**
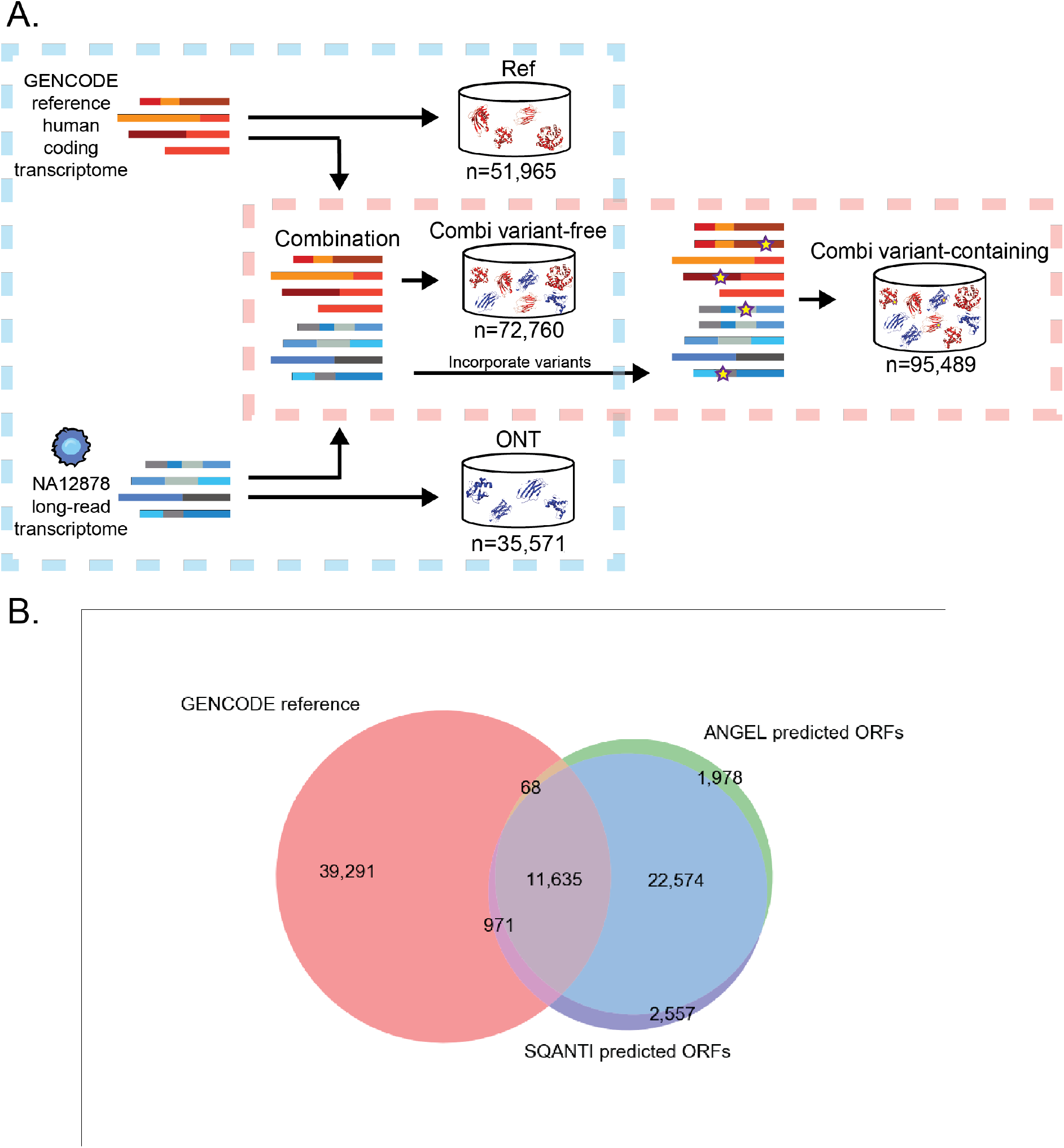
Creation of the search databases. (**A**) Three databases were made to make comparison between use of different sources of sequences. One with only translations of transcriptome sequences (ONT), one with only the reference proteome (GENCODE), and one with the union of the two. This comparison is denoted with a blue square. Variants from NA12878 were incorporated into the combination database from A and compared to the combination database without variants. This comparison is denoted with a red square. (B) The number of (predicted) ORF in the different sources used to construct the VF search database and their overlap. The sources included the GENCODE v29 reference ORFs and the predicted ORFs from ONT RNAseq. Two ORF prediction softwares (ANGEL and SQANTI) were used to determine candidate ORFs and the intersection was included in the final search database.

The Ref search database was made by filtering GENCODE v29 predicted ORFs for those that were complete (no 5’ or 3’ missingness). The ONT database was created using transcript structures provided by the NA12878 consortium (https://github.com/nanopore-wgs-consortium/NA12878/blob/master/RNA.md). The coordinates in the junction file (PSL format) provided were converted to BED with BEDOPS ^45^ and used to fetch the corresponding stretch of sequence from the GRCh38 genome with bedtools ^46^ getfasta. The exons were assembled using in-house scripts to form the full transcripts, and those that were non-identical to transcript sequences in GENCODE (“novel”) were then submitted to 2 ORF prediction software; ANGEL v2.4 (“dumb” ORF prediction on default settings) and SQANTI2 v2.7(https://github.com/Magdoll/SQANTI2). The translation of transcripts predicted by both prediction programs were added to the search database. ORFs from GENCODE were used for transcript sequences in ONT identical to transcript sequences in GENCODE.

The VF database was simply the union of the Ref and ONT databases. The VC database was created by first creating full-length coding sequences (CDS) with variants included by replacing reference nucleotides according to the VCF file per CDS fragment, for every CDS fragment. If only homozygous variant(s) were present in a CDS fragment, only one variant CDS fragment was generated. If a CDS contained at least one heterozygous variant, two variant CDS sequences were generated corresponding to the different alleles. Fragments were then assembled to full CDS. If a full CDS contained at least one CDS fragment with a heterozygous variant, two full CDS were generated corresponding to each allele. For those full CDS that contained at least one variant, the variant version(s) of the sequences replaced the non-variant versions in the VF database to create the VC database.

### Spectral search and post processing

Each run from Wu et al 2003 was first converted to the Mascot Generic Format(MGF) format using msconvert ^47^ with MS2 peak picking enabled. Each dataset was then searched against the four search databases described in the previous section, using *ionbot* version 0.5. Fixed and variable modifications were set according to the protocol in Wu et al. Open modification settings were enabled for all four runs, while open variant settings (for SAAV detection) were enabled for all runs except for on the VC database. Searches allowed for up to two missed cleavages. When parsing the search results, only spectra with an observed TMT6plex reporter ion 126.77 (corresponds to cell line NA12878) were retained.

Since sub-setting PSMs into groups such as variant peptides requires separate FDR correction ^31^, both VC and VF underwent a separate FDR correction for the variant peptide subset. Successful FDR correction requires the modeling of potential false positive peptide identifications using appropriate decoy peptides. In the case of variant peptides, this means a sufficient number of decoy variant peptide identifications must be present to accurately model the population of false positive peptides. Reversed sequences thus underwent the same processing steps as the true sequences in order to create the appropriate decoys. The distributions were checked for successful modeling (Figure S2).

A variant peptide list was created to compare with *ionbot* identifications from searches of the VC and VF. The list was created with an in-house Python script that performs an in-silico trypsin digest (allowing for up to 2 missed cleavages) with the pyteomics v 4.2 ^48^ package and checks per protein for peptides that differ by only one amino acid between the VF and VC database. I and L were treated as identical, and a potential variant peptide was disqualified if it appears in any other reference protein sequence.

*ionbot* identifications presumed to be variant peptides (and variant peptide decoys) underwent subset-specific FDR correction for both combination databases, but the exact subset of variant peptides differed between the two searches due to different assumptions. The assumption in the VF database is that variants in the genome are unknown, so all predicted variant peptides (and predicted variant decoy peptides) were pooled for FDR correction. In the VC database, only known variant peptides (and corresponding decoy peptides) are pooled for FDR correction. Q value calculation and cutoff (q < 0.01) were performed with an in-house python script (distribution can be seen in Figure S2). Retention time predictions were calculated with DeepLC ^49^. All scripts referred to in this manuscript can be found in the GitHub repository (https://github.com/cmbi/NA12878-saav-detection).

## Results

### Search database makeup

The main goal of this study is to evaluate the added value of transcriptomics data for SAAV identification in proteomics data. In this evaluation, SAAV identification with and without transcriptomics prior knowledge is compared for a state-of-the-art open search engine. To this end, we searched the NA12878 deep shotgun proteomics data set with four distinct search databases corresponding to two comparisons, as outlined in Figure 1A. The first comparison was between databases based on the Oxford Nanopore (ONT) long-read transcriptome, the GENCODE reference proteome (referred to as Ref) or the combination of the two (referred to as combi, Figure 1B). In this comparison, all searches were run with open modification settings that allow for one mutation in the peptide match. The second comparison was between a regular and an open variant search using databases that did and did not include NA12878 genome sequencing-derived variants, respectively. This comparison was performed for the combi databases only. The analysis with the variant-free combi database will be referred to as the VF method and the analysis with the variant-containing combi database will be referred to as the VC method. In this comparison, open modification search was enabled for both methods, but open variant search was only enabled in the VF method to allow for the detection of SAAVs. Open variant search is disabled in the VC method, because the variants were already incorporated in the VC search database.

### Adding the long-read transcriptome for the cell line does not contribute to additional peptide identifications in practice

Reliable peptide identification normally requires a comprehensive search database. We first investigated whether novel transcripts from long-read transcriptome sequencing would contribute to peptide identifications in the NA12878 shotgun proteomics data. The ONT database contained 35,248 full-length transcript sequences, 64% of which were novel. Although the combi database containing these novel predicted ORFs was 42% larger than the Ref database (Figure 1B), the number of unique peptides from these sequences made up a mere 2.3% of the search database (Figure 2, top panel). The addition of ONT-derived ORFs to the Ref ORFs thus translated to an only modest increase in the number of unique peptides in the search database (Figure 2, lower panel). A likely explanation for this is the fact that many of the novel ONT transcripts demonstrate high similarity to existing reference sequences. The sequences usually only differed in the length of the 3’ or 5’ UTR or the in use of alternative exon junctions rather than completely novel exons. The number of observed peptide matches to the novel ONT transcripts was even smaller: only 0.3% of unique peptides identified to the combi databases mapped exclusively to novel ONT transcripts. This indicates that the transcriptome database does not contribute significantly to the proteomic search results and suggests that alternative splicing and mRNA processing events do not contribute much to the diversity of the MS-detectable fraction of the proteome.

**Figure 2.**
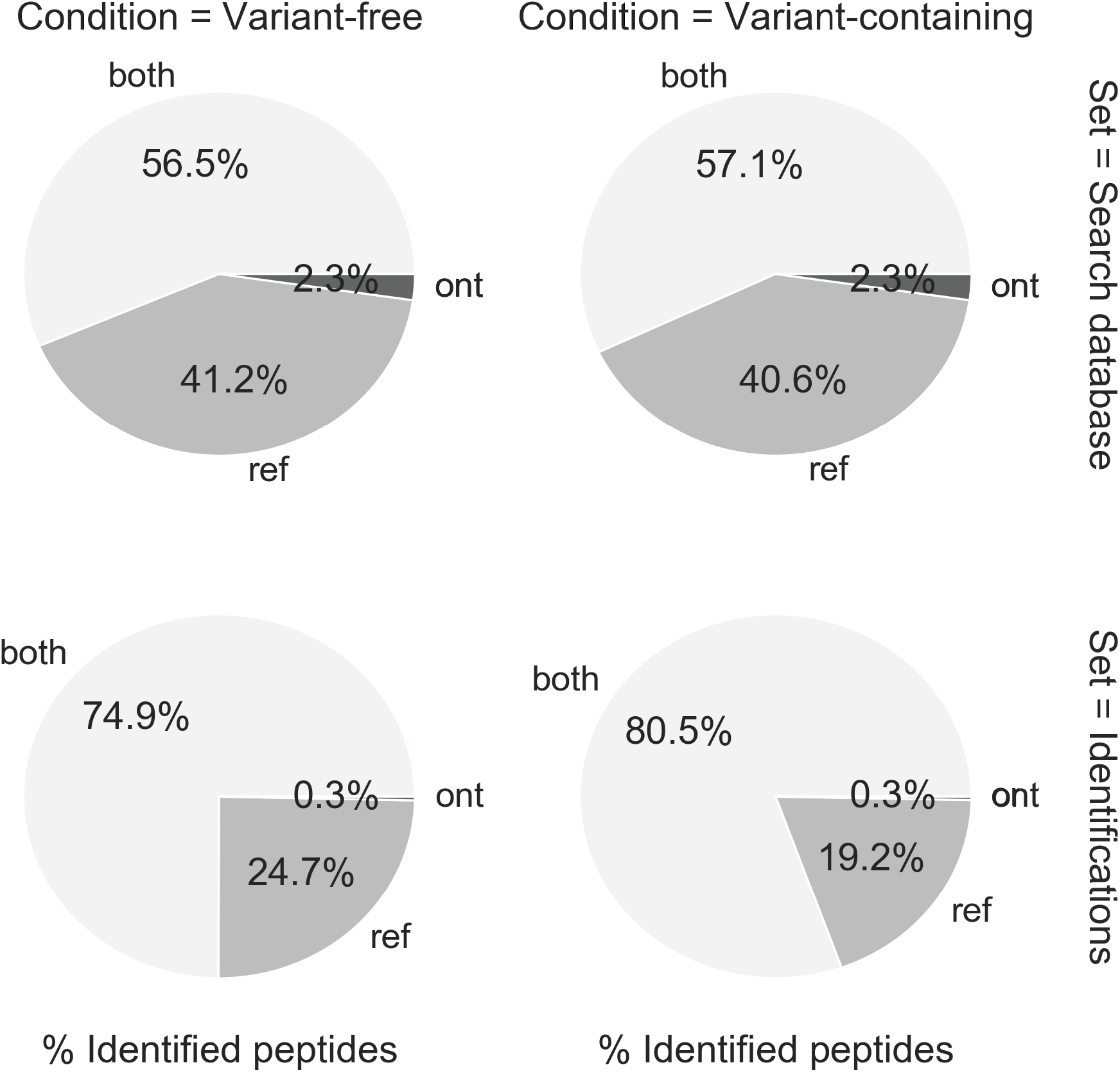
Detectible peptides per method. Theoretical (upper pie charts) and observed (lower pie charts) proportions of peptides when searching against VC (right) or VF (left) search databases. This shows percentages of matched peptides attributed only to GENCODE proteins, only ONT proteins, and those that match to proteins in both databases.

Aside from the contributions from the ONT-only sequences, it is also interesting to investigate protein identifications that were not found in the ONT transcriptome. While these should theoretically not be present, roughly 20% of identified peptides are exclusively matched with the ENCODE transcripts (Figure 2). As expected, this percentage is smaller than the 42% of peptides in the search database that are exclusive to GENCODE transcripts, but still a significant fraction. This suggests that it is best to still use a reference transcript database, even if there is full transcriptome sequencing data available.

### Variant-containing method allows detection of many more genome-supported variant peptides

We subsequently studied the effect of the inclusion of sample-specific variants in the search database. In the VF method, the data is analyzed with an open variant search, thus letting the search engine predict single amino acid substitutions. This is in contrast to the VC method, where no variants are predicted and only genetically supported variant are present in the search database. We detected 461 variant peptides by the VC method and 62 by the VF method, with 59 overlapping between the two methods (Figure 3A). The greater majority of variant peptides that were detected by the VF method only (n=1,805), were not supported by the genome and are likely false positives (Figure 3B). In addition, one third of variant peptide matches that appeared to be supported by the genome, actually contained an incorrect amino acid substitution. Thus, the inclusion of variant peptides derived from personal genomes in search databases is far superior to the use of a variant free database combined with an open variant search.

**Figure 3.**
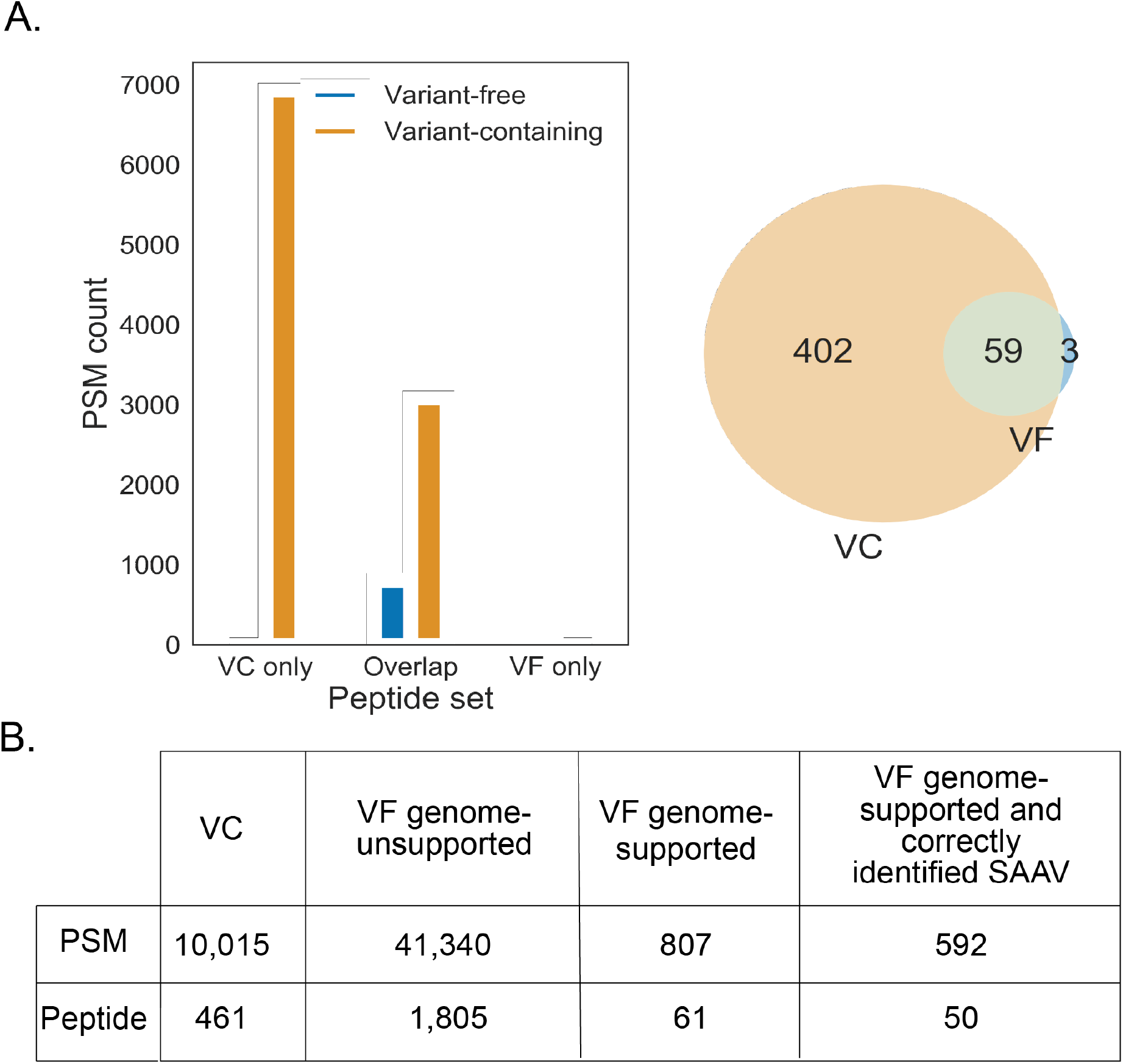
Detection of variant peptides using (combination) VF and VC databases. (A) variant PSMs (left) and unique peptides (right) attributed to genome-supported variant peptides. (B) PSM and peptide counts found by each method.

### Detectible variant peptides have attributes that differ from expected variant peptides

Out of the 34,968 peptides in the genome-supported variant peptide list, only 462 were detected by either or both the VC and VF methods (Figure 4A). They are not a random sample of all possible variant peptides. Namely, some variant peptides are easier to detect than others depending on their abundance and/or properties, and that differs even between methods. For instance, the VF method tends to find longer variant peptides (in a range of 16-27 aa) and misses the shorter variant peptides (Figure 4B). This highlights the larger amount of ambiguity in variant peptide identification proportional to the lower number of peaks in the spectra. The VC method does not suffer from this ambiguity and allows for detection of a wider range of variant peptide lengths than variant-free, especially shorter variant peptides (p=0.0017 K2 samp). While there is a bias in variant peptide length, we did not find clear evidence that the position of the variant within the peptide affects detection of the variant peptide in either of the methods. In addition, the amino acid substitution itself affects detectability, since the corresponding mass shift in the MS/MS spectrum needs to be separated from noise or similar mass shifts corresponding to other modifications in order to be identified. There are some predefined limitations to SAAV detection with the VF method that lead to certain amino acid substitutions getting detected less than expected (Figure 4C). Amino acids on which there are fixed modifications can’t have variants in the open variant search, meaning substitutions at K and C are not detected. Substitutions affecting the trypsin digest, such as those involving R, can also not be detected.

**Figure 4.**
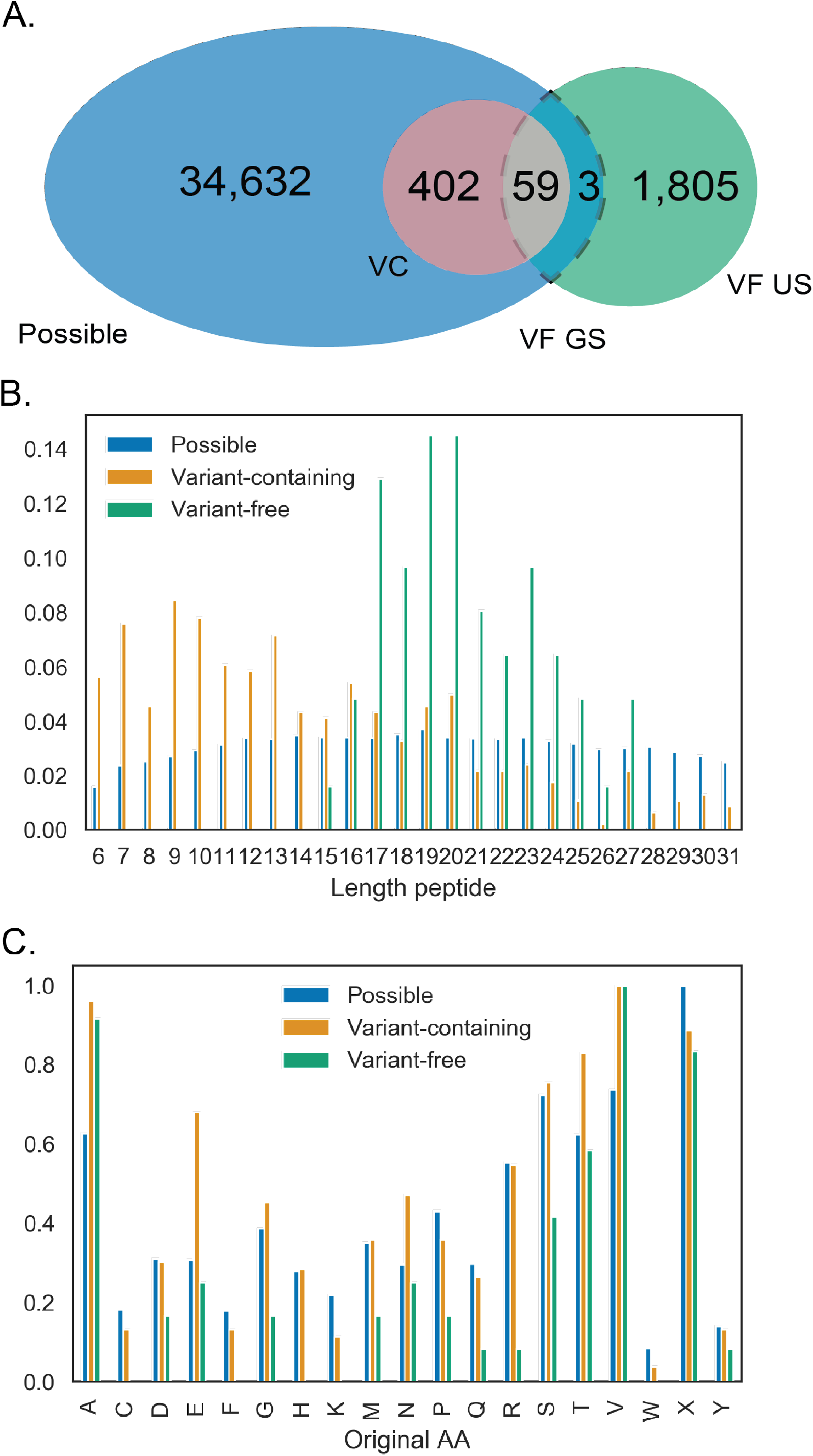
Properties of detected variants compared to expected. (A) The groups of variant peptides being compared. Each circle, including all overlaps, are being compared to each other. (B) Length distribution differences between detected variant peptides by the different variant detection methods. (C) Normalized (divided by max) frequency of variation per original (reference) amino acid.

### Erroneous variant peptide identifications are difficult to discern from true variant peptide identifications

The misidentifications from the open variant-free approach can be separated into false negatives and false positives. False negative identification is where the VC method identifies variant peptides, but those same spectra are identified by the VF method as non-variant peptides. False positive misidentification is where the VF method identified variant peptides that were not supported by the genome.

There were 402 unique false negative peptides observed (Figure 5A). These false negatives peptides were classified as variant peptides by VC method but not by VF, although they were contained in the VF search space. Identifying causes of false negatives requires investigation of how the VC peptides were identified with the VF method. There was no particular length peptide that was mis-identified more than others in general, despite the difference in detectible peptide length (Figure S3). The peptide identifications were similar between the VF and VC methods. In general, length correlated highly between the identifications of the two methods (R^2^=0.9071, p=0). When comparing individual peptide identifications per method for mismatches and length difference, the largest source of error was a 1 aa length difference. Nonvariant peptides with a 1 aa length difference from the variant peptide were being identified instead of the correct variant peptide in >30% of the false negatives (Figure S3). Another possible source of false negative errors that was investigated is SAAVs being mistaken for unexpected post-translational modifications. In the false negative set, this did not appear to be an issue. The false negative VF identifications had approximately the same rate of unexpected PTMs (Figure S3).

**Figure 5.**
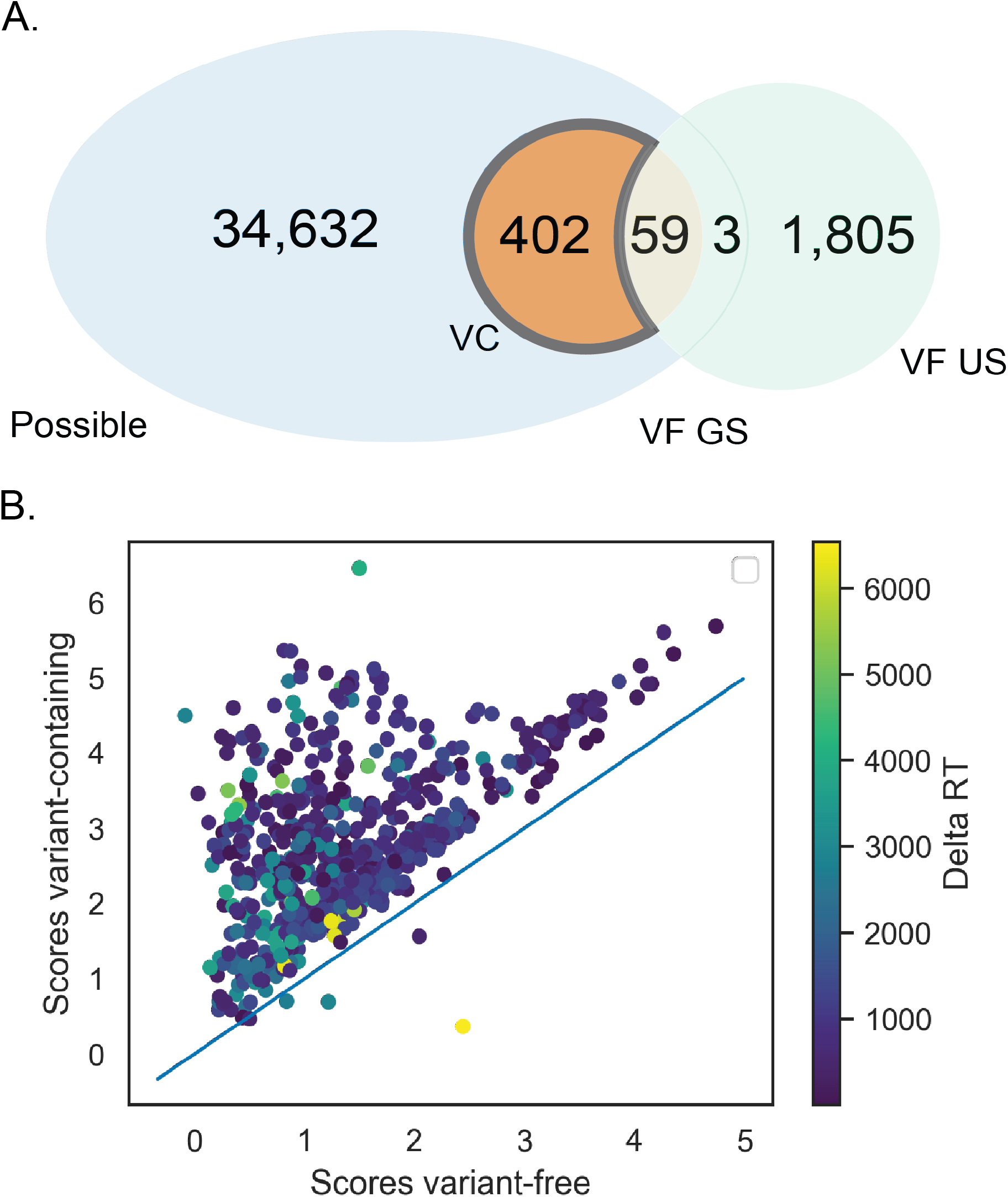
False negative variant misidentifications. (A) Investigation of causes of mis-identification of peptides in the variant-free set. (B) Scores of those mis-identified peptides in VF vs VC set. Each point corresponds to one false negative variant peptide. Percolator PSM score is used. Color corresponds to delta retention time.

To further understand how false negatives could occur, we compared the peptide matching scores of the false negative spectra for the VF and VC search methods (Figure 5B). Higher scores indicate higher confidence in assignment of spectra. VC scores for false negative peptides were generally higher than the VF scores (mean score ratio VC/VF = 1.31). However, a large fraction of the false negatives received comparable scores in the VC and VF search methods. This could indicate a ranking problem: the variant peptide received a score equal to another peptide, to which the peptide spectrum was ultimately assigned. Delta retention time can often be a useful independent validator when score disagrees between the different search methods. Despite high retention time discrepancies in this particular data set, observed retention time aligns relatively well with predicted retention time for those spectra that received higher scores in VC.

The genome-supported variants are a tiny fraction of the high confidence variant peptide predictions from the VF database, indicating a high false positive rate (Figure 6A). We investigated whether there are distinguishing features between genome-supported and genome-unsupported variants. Reassuringly, scores of true positives were slightly higher than false positives (Figure 6B, p=1.34e-26, ANOVA).

**Figure 6.**
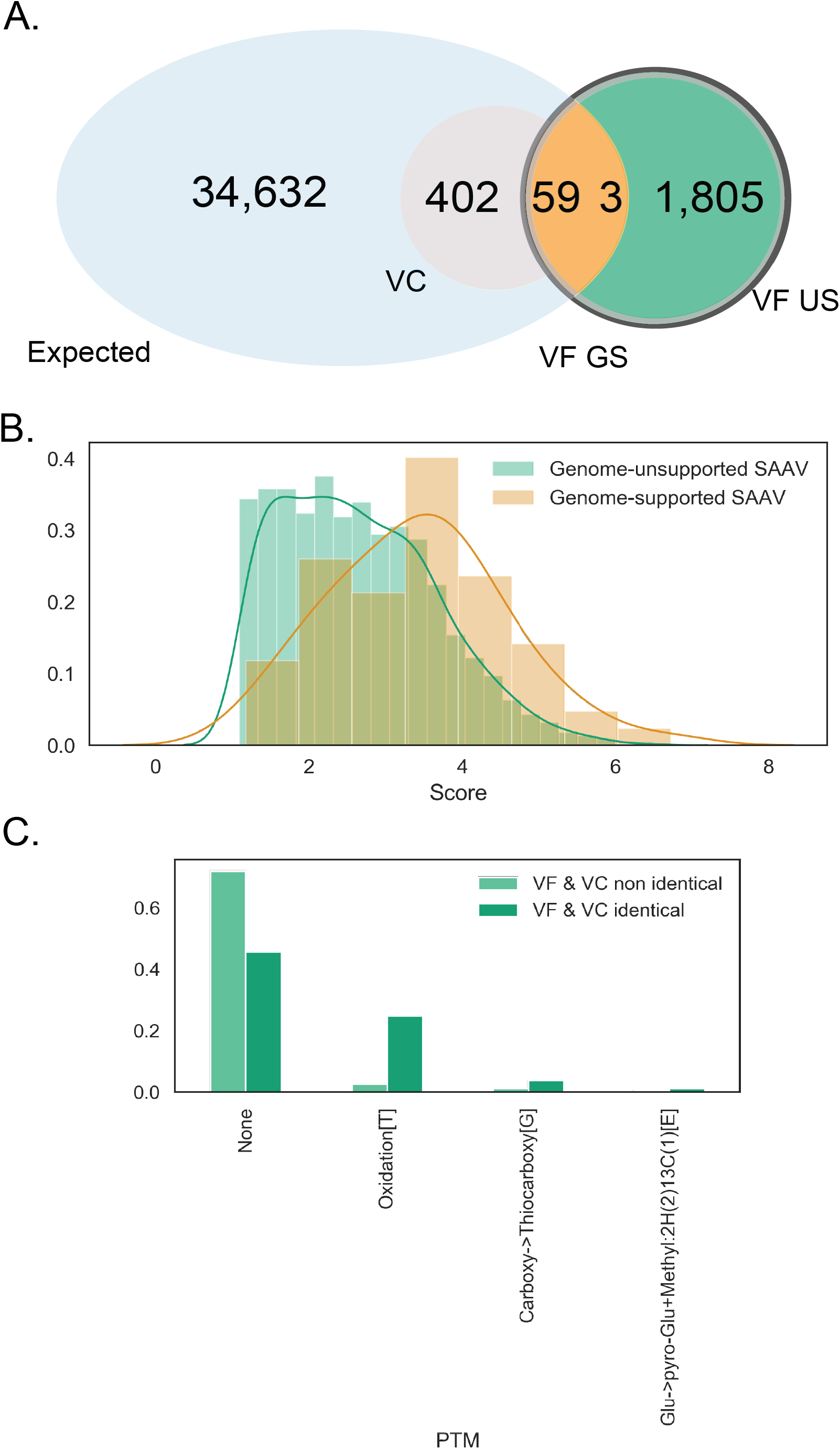
False positive misidentifications. (A) False positive misidentifications are genome-unsupported (US) variants predicted by the variant-free method (VF). The venn diagram highlights the subset of variants that are being investigated in this figure. These 2,998 variants were predicted by ionbot to be variant peptides, but were not found with the variant containing set. All but 7 were variants unsupported by genome information. (B) Relative score distributions between genome supported vs unsupported variants in the variant-free set. (C) Unexpected modifications by the VC set corresponding to all ‘false positive’ predicted variant PSMs in the VF set.

A closer inspection of genome-unsupported variants reveals potential sources of confusion for variant prediction algorithms, leading to false positive identifications. There was a high level of concordance of peptides matched to these spectra in general. Two thirds of spectra that corresponded to genome-unsupported variant peptide identifications by VF had the same base peptide identifications in both the VF and VC searches. Mass shifts predicted to be SAAV in VF were commonly predicted to be ‘unexpected’ PTMs by the VC method (Figure 6C). A common PTM mistaken as a SAAV in VF was threonine oxidation, but many PTMs contributed to this mix-up. There was no clear trend to the identification errors, underlining the difficulty of correctly classifying minor mass shifts corresponding to PTMs and SAAVs.

### Evaluation of the variant peptides’ SNPs of origin

The detection of variant peptides is ultimately a means to understanding which single nucleotide variants (SNVs) are expressed on the protein level. By incorporating SNVs into predicted ORFs, we ended up with a theoretical set of 34,968 variant peptides originating from 9,298 SNVs from all chromosomes, of which 5,989 are heterozygous variants.

In the case of a heterozygous variant, both variant peptides and their reference counterparts can be identified in some ratio. A ratio different from 0.5 may be indicative of preferred expression of one of the alleles on the protein level, otherwise known as ASPE (allele specific protein expression). Presence and magnitude of ASPE is potentially key information that can be used to understand biological mechanisms. However, technical biases of search methodology may invalidate potential findings by distorting these ratios. For the VF method, the reference peptide was identified more frequently than the variant peptide (p=0.013, oneway ANOVA). The opposite was true for the VC method.

Homozygous variants can be used as a type of control to understand the bias in search methods, since we know that only one of the two alleles can be expressed. In case of homozygous variants, the variant peptide is expected to be present in all cases – with no reference counterpart. This was observed for the VC but not for the VF method (Figure 7A). Thus, without prior information about zygosity, the VF method tends to be conservative in identifying SAAV peptides, resulting in a higher likelihood of the reference peptide than its variant counterpart.

**Figure 7.**
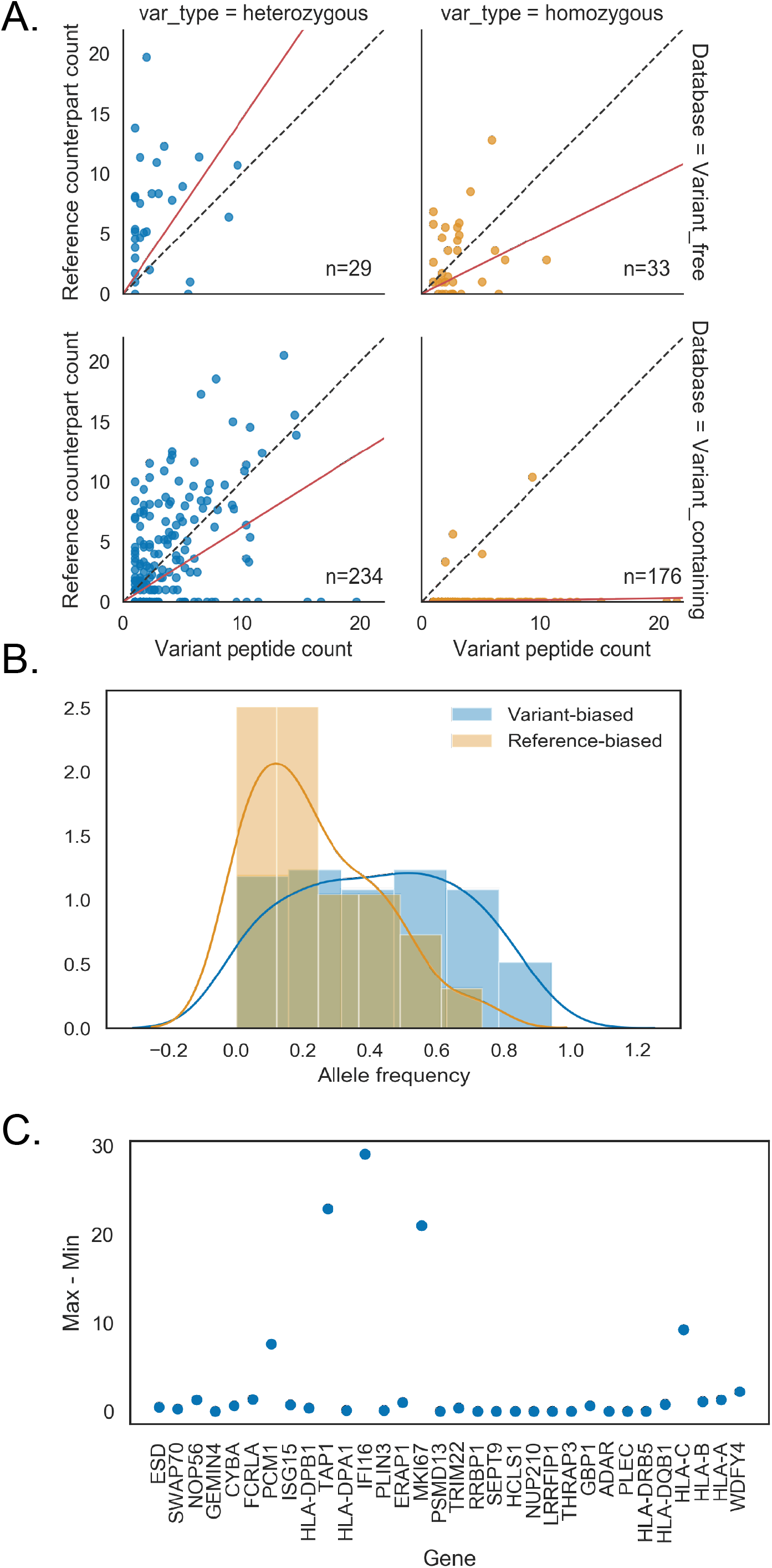
Underlying SNPs detected on the protein level. (A) Variant peptide abundance vs reference counterpart split by zygosity and search database, square root transformed. (B) Separating heterozygous variants in the variant-containing database by whether more variant peptide was found (variant-biased) or more of the reference counterpart was found (reference-biased) revealed differences in allele frequency distributions. (C) Ratio variability of genes with 2 or more variant peptides. Ratio is defined by the variant counterpart abundance divided by variant peptide abundance. Y axis shows max – min per gene.

It is evident that some variant peptides were observed much more often than their reference counterparts or vice versa. The VC heterozygous variant peptide identifications should not suffer from the technical reference bias and allow for detection of allele specific expression on the protein level. The VC-detected heterozygous variants were divided in two groups; one group with more counts for the reference peptide (reference-biased, N=78) and one group with more counts for the alternative peptide (alternative-biased, N=123). The two groups demonstrated a clear and significant difference in population allele frequency (p=6.45e-08. Figure 7B). Those with lower allele frequencies displayed a stronger reference bias. This could be explained by the fact that rare variants in coding regions have a higher likelihood of causing undesirable effects on the resulting protein. Any deleterious effects resulting from the variant on protein stability would be visible as depletion of the alternative allele.

One significant subgroup of heterozygous variants was particularly biased towards the alternative allele. Fourty-four out of 183 variant peptides supported by more than two PSMs did not have any detected reference counterparts. One third of these variants had a substitution involving arginine or lysine (tryptic cleavage sites). One gene, HLA-DBQ1, had two alternative alleles instead of one reference and one alternative. In general, the score distribution for these highly biased group was lower than the score distribution for all VC detected variant peptides. The allele frequencies of this group were not different to those of the overall alternative-biased group (p=0.5, ANOVA). There was also no correlation between the RNA expression of these genes to the variant peptide expression (R^2^=0.01). Also, a comparison the list of genes displaying ASE on the RNA level from ^42^ to the heterozygous genes with variant peptides detected on the protein level yielded negligible overlap (2 genes).

A total of 33 genes were detected through two or more unique variant peptides. For variant peptides within a gene, the reference peptide to variant peptide ratio should be consistent, unless there are different protein isoforms as a consequence of alternative splicing. This was the case for the majority of genes with multiple variant peptides belonging to the same gene (Figure 7C). Five of these genes were represented by multiple variant peptides with inconsistent ratios. HLA-C, IFI16 and MKI67 had peptides matching to non-identical (sets of) isoforms within the gene. PCM1 had peptides matched to 24 isoforms. That is four times the average number of isoforms matched by a variant peptide in the VC search. Thus, inconsistent variant to reference peptide ratios within a gene can generally by attributed to differing abundances of protein isoforms.

## Discussion

Here, we have carried out an investigation of the effects of proteogenomic additions to a proteomics search database. To this end, we compared a typical proteomics approach to a purely proteomics method utilizing state-of-the-art open search. We observed that the addition of transcriptomic sequences to the search database did not have significant effects on the overall peptide identification rate. There was a roughly equal number PSMs from the three databases, despite the long-read transcriptome search database being 40% smaller than that of the union of it and the reference. At the same time, the matches to reference-only sequences in the combination database imply that >20% of peptide identifications are missed. This suggests a large portion of false identifications when using a database comprised of only ONT sequences.

The fact that around a quarter of peptide identifications cannot be attributed to the transcriptomics data is rather surprising. There are a couple possible explanations. Using transcriptomics data from different cells than the proteomics data (different labs and different year) will unavoidably cause some discrepancies ^50^. This could also be attributed to protein stability in the cell, as proteins are detectable for some time after RNA have already been degraded ^51^. Also notable is the fact that including the transcriptome sequences did not seem to add significantly to the peptide detections; the proportion of novel peptides found was lower than the proportion of novel transcripts found. As this cell line/organism is so well studied, it is likely that the vast majority of present proteins have already been characterized. For other cell types and organisms with more novel transcripts, adding (full length) transcriptomes may lead to more peptide identifications.

Two different search methods were used to identify non-reference peptides derived from SNVs: a proteogenomics approach, in which all variants known from the genome sequence were added to the search database, and an ‘open variant search’, where only reference peptides were included in the search database and one amino acid differences were allowed by the search engine. The proteogenomics approach was clearly superior, as it detected 7-times more variant peptides, whereas the open variant search suffered from many false positive identifications that were not supported by the genome sequence, and from large numbers of false negatives. Nevertheless, also the proteogenomics search method detected only a minor fraction of the variant peptides predicted to be present in the genome. It has been estimated before that maximum ~70% of variants in protein coding regions are theoretically detectible in an ideal shotgun proteomics experiment considering peptide lengths 7-40 aa ^52^. The number of variants found with a proteogenomics method in practice is much lower, depending on method details. Some studies either use a statistically dubious ‘multi-tier’ method ^53, 54^ or skip FDR sub-setting altogether ^55^ and report the number of variants detected to be in the region of 10%. We detect only 1% of the theoretically present variant peptides, despite the ~4M spectra present in this dataset, making it one of the deepest proteomics datasets currently available. This is partly due to the careful control of FDRs in our study. Also other conservative efforts to detect variant peptides using FDR sub-setting or targeted proteomics validation detect <1% of all theoretically present variant peptides ^23,54,56^.

While open search lags behind the proteogenomics approach for the moment, it has promise. Algorithms are being continuously improved to better differentiate signal from noise, which will reduce the false positives and false negatives in variant peptide detection ^57^. There are several upcoming methodologies to further refine the open search to increase accuracy, either adding to existing peptide identification tools or standalone with promising results such as Open-pfind ^58^, TagGraph ^39^, MSFragger ^59^, Crystal-C ^60^. There are considerable challenges still to face in their detection, particularly in noise/signal differentiation. This is especially complicated as variants often co-occur with other PTMs such as phosphorylation ^30,54^. Current detection methods including *ionbot* cannot handle the complexity of two modifications on one site. However, deep neural networks show great promise with difficult peptide identifications ^61^. Using methods of machine learning along with orthogonal information such as peptide retention time should result in significant improvements in open search ^62^. This in combination with rapidly improving data-independent acquisition removes detection limitations of low-abundance or otherwise difficult to detect peptides ^63^, which is currently a considerable hurdle in SAAV peptide detection ^55^. Including open search is clearly useful and bound to get more accurate. This study used *ionbot* as the sole predictor of unexpected modifications/SAAVs, and comparison between identification tools was difficult as no other identification software tested reported the precise reporter ions per matched spectra (to be able to separate TMT tags corresponding to different cell lines). A study to compare methods given these updates is certainly warranted and ensemble methods may eventually be used to even more accurately predict these unexpected modifications/SAAVs.

One important implication of correctly detecting SAAVs is the ability to observe allele specific expression on the protein level. A targeted proteomics approach has recently been described to study ASPE (allele specific protein expression) with high confidence ^64^. It found no correlation between RNA and protein level ASE for the few variants studied, highlighting the utility of having higher throughput methods to study this phenomenon. One simple way to measure ASPE when using a proteogenomics approach is by comparing the spectral counts for the SAAV and its reference counterpart, since a reference counterpart usually has equal detectability by MS/MS ^52^. Here we found low correlation between the abundance of the variant and reference counterparts regardless of VF or VC method. This is potentially indicative for a high level of ASPE. In contrast, ^54^ demonstrated a high correlation between variant and reference peptides. This may be attributed to the low stringency associated with using the multitier search strategy for SAAV detection. We found no correlation between ASE and ASPE was found in this study which is consistent with the findings of Shi et al.

## Conclusions

Our study provides guidance for the detection of variant peptides that shape the personal proteome. While personal genomes currently seem indispensable for the characterization of personal proteomes, new computational and analytical tools and new file formats to accommodate personal proteome information will allow us to get the fullest picture possible of the individual proteome, even without personal genome information.

## Supporting information

All supplementary

## Acknowledgements

This work is funded by the Radboud Institute for Molecular Life Sciences.

## SUPPORTING INFORMATION

**Figure S1.**
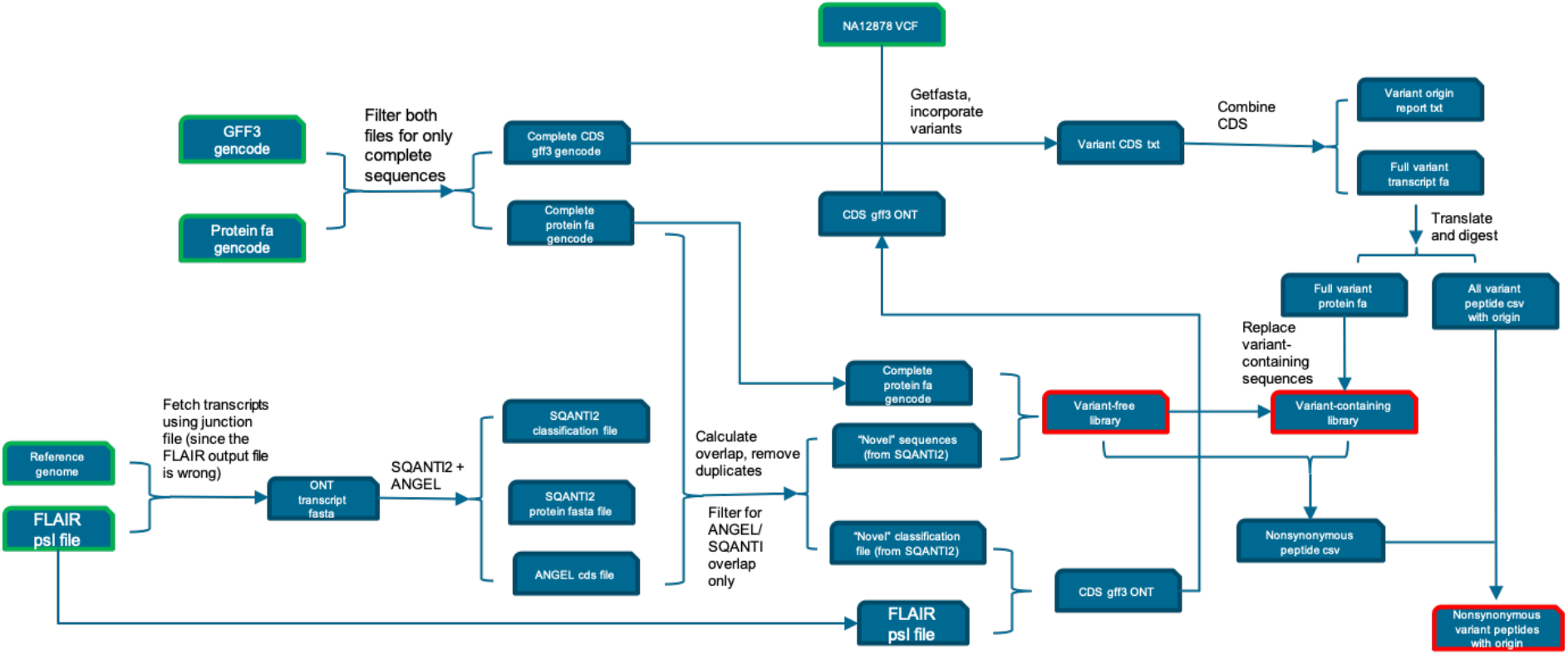
Detailed workflow schematic.

**Figure S2.**
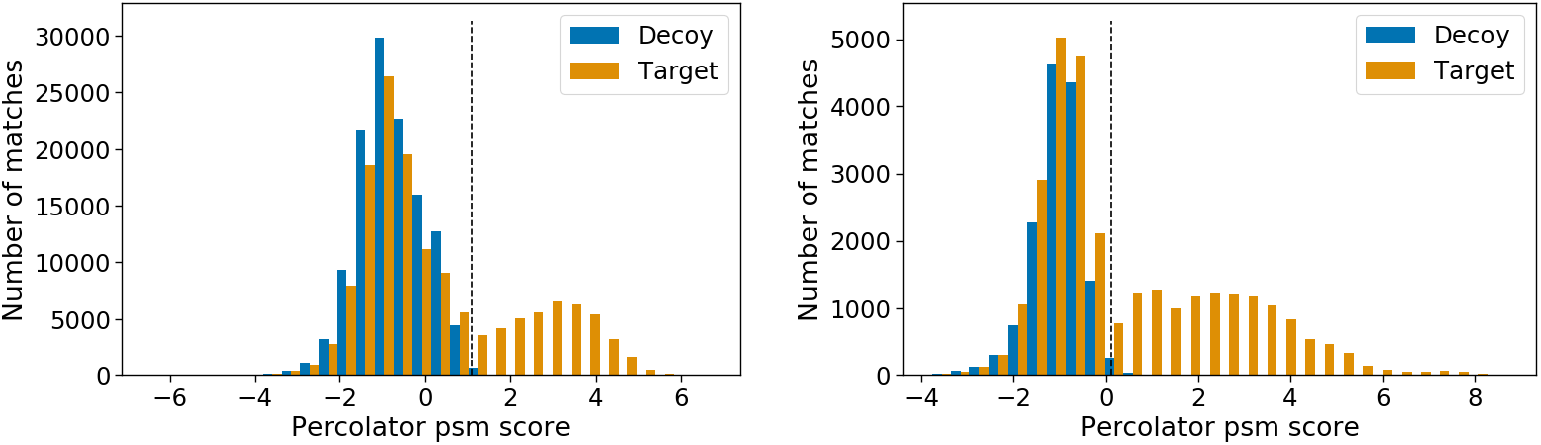
Distribution of target and decoy variant peptides. Variant-containing distribution is on the left, and variant-free is on the right. Separation was made at the dotted line (q<0.01).

**Figure S3.**
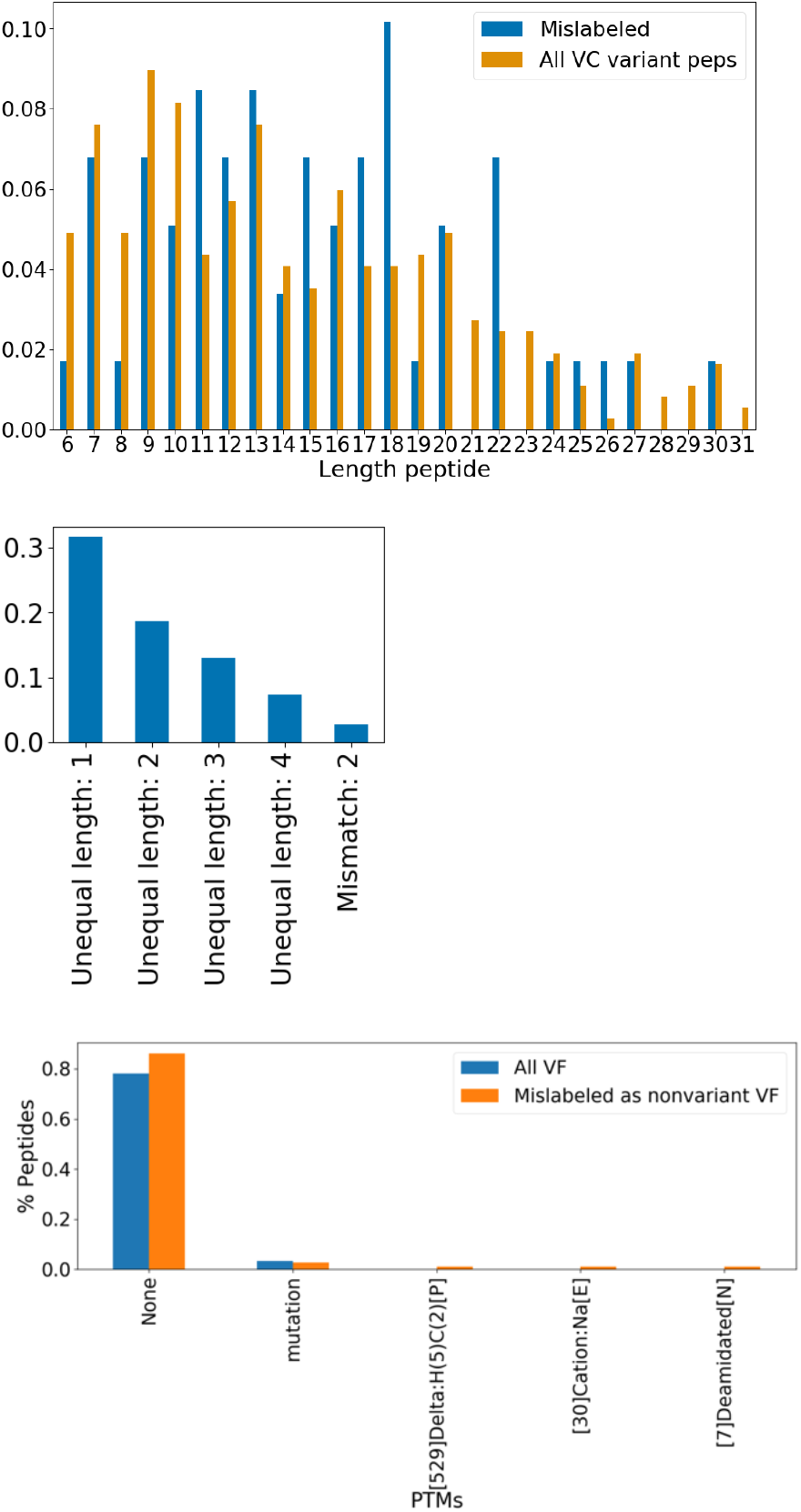
Investigation of false negative (‘mislabeled’) identifications by *ionbot.* Top figure shows the density of mislabeled peptides per length, as compared to lengths of all variant peptides identified by the VC method. Middle figure shows the 5 most common causes of misidentification of variant peptides by *ionbot*. Bottom figure shows unexpected modifications of the false negatives versus the unexpected modifications by all VF identifications. Unlabeled y axises refer to density.

**Table S1.**
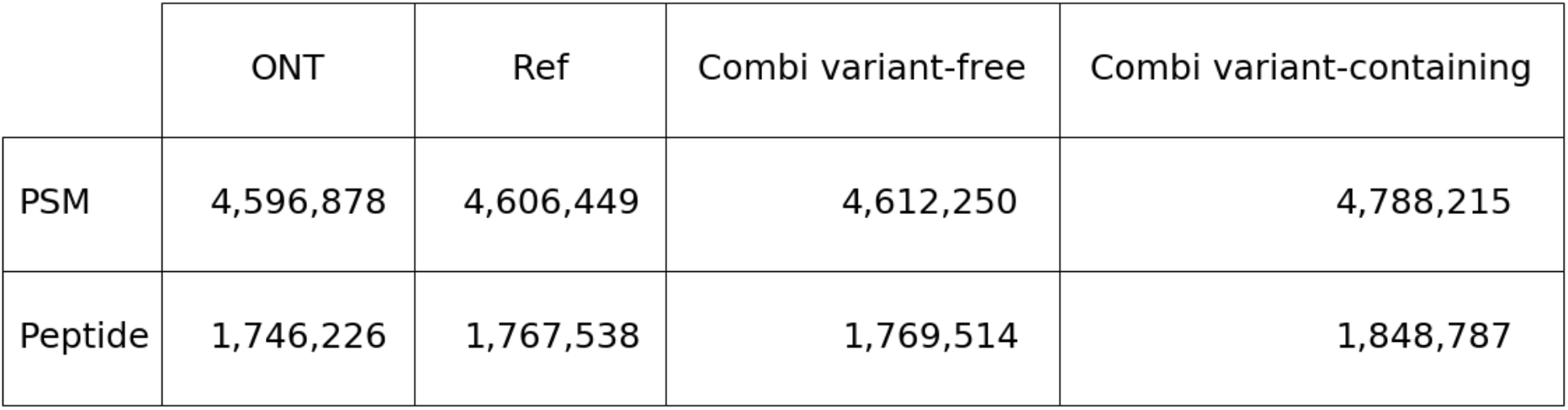
Absolute numbers of PSMs and peptides detected per method.

