## Supplementary material for "The Personalized Proteome: Comparing Proteogenomics and Open Variant Search Approaches for Single Amino Acid Variant Detection": All supplementary

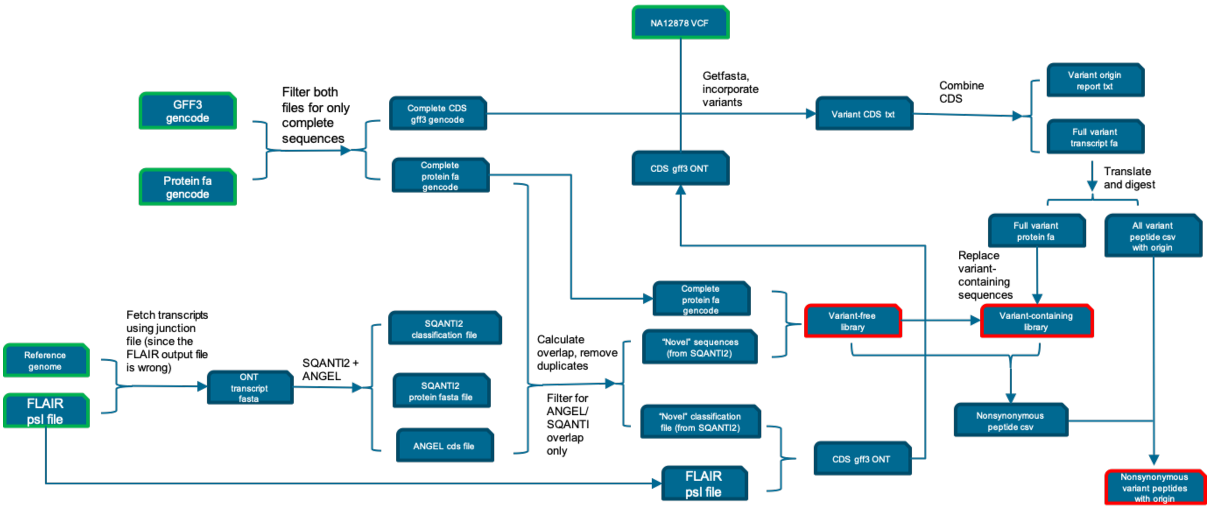


Figure S1. Detailed workflow schematic.

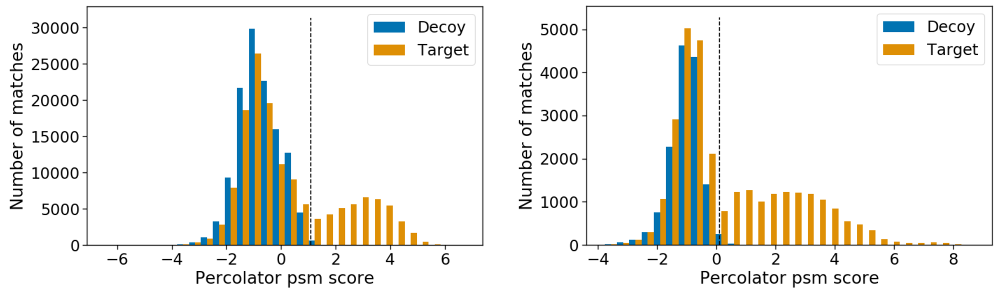


Figure S2. Distribution of target and decoy variant peptides. Variant-containing distribution is on the left, and variant-free is on the right. Separation was made at the dotted line (q<0.01).

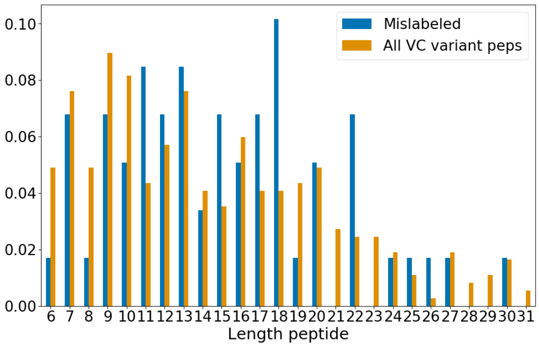


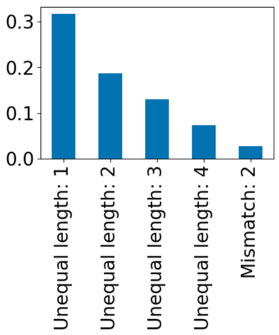

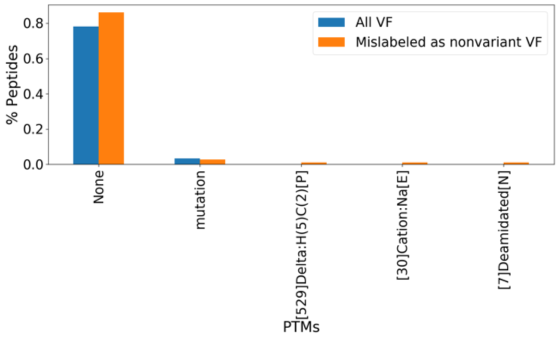


Figure S3. Investigation of false negative ('mislabeled') identifications by *ionbot*. Top figure shows the density of mislabeled peptides per length, as compared to lengths of all variant peptides identified by the VC method. Middle figure shows the 5 most common causes of misidentification of variant peptides by ionbot. Bottom figure shows unexpected modifications of the false negatives versus the unexpected modifications by all VF identifications. Unlabeled y axises refer to density.

| | ONT | Ref | Combi variant-free | Combi variant-containing |
|---|---|---|---|---|
| PSM | 4,596,878 | 4,606,449 | 4,612,250 | 4,788,215 |
| Peptide | 1,746,226 | 1,767,538 | 1,769,514 | 1,848,787 |

Table S1. Absolute numbers of PSMs and peptides detected per method.
